# Near-complete telomere-to-telomere *de novo* genome assemblies of Egyptian clover (*Trifolium alexandrinum*)

**DOI:** 10.1101/2024.07.04.602140

**Authors:** Mitsuhiko P. Sato, Ramadan A. Arafa, Mohamed Rakha, Amero A. Emeran, Sachiko Isobe, Kenta Shirasawa

## Abstract

Egyptian clover (*Trifolium alexandrinum* L.), also known as berseem clover, is an important forage crop to semi-arid conditions that was domesticated in ancient Egypt and introduced and well adapted to numerous countries. Despite its agricultural importance, genomic research on Egyptian clover has been limited to developing efficient modern breeding programs. In the present study, we constructed near-complete telomere-to-telomere-level genome assemblies for two Egyptian clover cultivars, Helaly and Fahl. Initial assemblies were established by using highly-fidelity long-read technology. To extend sequence contiguity, we developed a gap-targeted sequencing (GAP-Seq) method, in which contig ends are targeted for sequencing to obtain long reads bridging two contigs. The total length of the resultant chromosome-level assemblies was 547.7 Mb for Helaly and 536.3 Mb for Fahl. These differences in sequence length can be attributed to the expansion of DNA transposons. Population genomic analysis using single-nucleotide polymorphisms revealed 38 highly conserved genomic regions within Helaly. Growth- and stress response-associated gene ontologies were enriched in the 38 regions, indicating that these genes may determine the unique characteristics of Helaly. Comprehensive genomic resources can provide valuable insights into genetic improvements in Egyptian clover and legume genomics.

## Introduction

Forage crops are essential for livestock production. Clover, which encompasses a range of *Trifolium* species, is an important forage crop (Graham and Vance, 2003). The genus *Trifolium* is a member of the cool-season annual legume family and is genetically similar to the genus *Medicago* (Choi *et al*., 2004; Ellison *et al*., 2006). Among *Trifolium*, red clover (*T. pratense*) and white clover (*T. repens*) are popular species for forage production worldwide (Kjærgaard, 2003; Riday, 2010). Subterranean clover (*T. subterraneum*) is resilient to poor-quality soil where other clovers cannot survive (Nichols *et al*., 2013).

Egyptian clover (*T. alexandrinum*), or berseem clover, originated in the eastern Mediterranean region and was domesticated in ancient Egypt (Muhammad *et al*., 2014). Egyptian clover was introduced to numerous countries during the late nineteenth and early twentieth centuries (Muhammad *et al*., 2014). The primary targets of Egyptian clover breeding programs are high biomass production, ease of cultivation, and substantial nitrogen fixation capacity (Muhammad *et al*., 2014). Since this species is allogamous in nature, like red and white clovers, a mass selection method is usually applied in the breeding programs, resulting in the maintenance of genetic diversity within a cultivar to avoid inbreeding depression by self-crossings (Abdalla and Abd El-Naby Zeinab, 2012). Genomics is a promising tool for accelerating breeding programs by playing a crucial role in the rapid development of new varieties and ensuring resilience and adaptability in changing climates (Thudi *et al*., 2021). Although genome information on red clover, white clover, and subterranean clover is publicly available (Sato *et al*., 2005; Isobe *et al*., 2012; Bickhart *et al*., 2022; Santangelo *et al*., 2023; Shirasawa *et al*., 2023), no genome data for Egyptian clover have been reported thus far, except for the chromosome number of 2n = 2x = 16 (Wexelsen, 1928).

Genomic information is required to facilitate breeding programs for Egyptian clover with complex genetic backgrounds. The widely cultivated Egyptian clover cultivars in Egypt belong to the ‘Meskawi’ botanical group (multi-cut type), which can regrow after being cut and thus is capable of delivering multiple cuts during the growing season. Being a basal-branching type, ‘Meskawi’ is characterized by the aggregation of buds in the basal crown area, which is left after cutting to allow for regrowth (El-Naby *et al*., 2014). Helaly cultivar, which belongs to Meskawi realized depending on forage production vigor. On the other hand, single-cut ‘Fahl’ Egyptian clover is another botanical group of clover that lacks regrowth ability. As a stem-branching type, buds are vertically distributed along the main stem and, thus, removed by taking the first and only cut from the crop (Vijay *et al*., 2017).

Advances in high-throughput sequencing technologies, such as the use of ultra-long read sequences, co-barcoding of long DNA fragments, Hi-C, and optical mapping methods, have accelerated the determination of high-quality novel genome sequences at telomere-to-telomere (T2T) levels (Nurk *et al*., 2022; Kurokochi *et al*., 2023). Several plant genomes have been sequenced at the T2T level, including those with large genome sizes or tetraploid structures (Sato *et al*., 2023). However, establishment of T2T- or chromosome-level genome assemblies using advanced technologies is not always possible. A combination of long reads and target sequencing would be useful for extending sequence contiguity.

In this study, we presented the near-complete T2T-level genome sequences for two cultivars, Helaly for multi-cut type and Fahl for single-cut type. Genetic diversity within and between cultivars was evaluated on the basis of genome sequence data. These findings provide genetic and genomics context for Egyptian clover, paving the way for future breeding programs for Egyptian clover and other related species.

## Materials and Methods

### Plant materials

Two *T. alexandrinum* cultivars, Helaly and Fahl, were used in this study. Both cultivars were obtained from Agricultural Research Center, Giza, Egypt. Fahl is commonly used for single cut as it has poor regeneration ability. Helaly is commonly used in north of Egypt and can give 5-6 cuts of fodder, and has good regeneration ability after cutting (Abdel-Fatah and Bakheit, 2019).

### *De novo* genome sequencing and assembly

Leaf samples were collected from a single plant of each cultivar and subjected to high-molecular-weight genomic DNA extraction using a Genomic-tip 500/G (Qiagen, Hilden, Germany). Short-read sequence data were obtained to estimate the genome sizes of the Helaly and Fahl cultivars. The genome sequence libraries were constructed with Swift 2S® Turbo Flexible DNA Library Kit (Swift Biosciences, MI, USA) and sequenced on the DNBSEQ-G400 (MGI Tech, Shenzhen, China). Generated short-read sequences were processed by trimming low-quality bases (quality value < 10) and adapter sequences (AGATCGGAAGAGC) using PRINSEQ and fastx_clipper from the FASTX-Toolkit, respectively. The genome sizes of both cultivars were estimated using *k*-mer distribution analysis (*k* = 21) with Jellyfish software (v.2.3.0) (Marçais and Kingsford, 2011).

For *de novo* genome assembly, the high-molecular-weight genomic DNAs were sheared with a g-TUBE (Covaris, MA, USA) by centrifugation at 1,600 × g, and long-read sequencing libraries were prepared using the SMRTbell Express Template Prep Kit v2 with a barcoded overhang adapter kit (PacBio, CA, USA) for multiplexing. The resulting libraries were separated using BluePippin (Sage Science, MA, USA) to remove short DNA fragments (<20kb). Sequence data was obtained using the Sequel II system (PacBio), demultiplexed, and converted into HiFi reads using SMRT Link pipeline (PacBio). The HiFi reads were assembled in contig sequences using Hifiasm version 0.19.5 (Cheng *et al*., 2021), in which the parameter that removed 20 bases from both ends of the reads was set.

The obtained contig sequences were scaffolded into pseudomolecule sequences using a strategy involving another type of long-read sequence and alignment between the cultivars and among closely related species (Figure 1). To link the contigs with another long-read sequencing technology, libraries were prepared using the Ligation Sequencing Kit V14 (Oxford Nanopore Technologies, Oxford, UK) and sequenced using a flow cell (R10.4.1) with GridION (Oxford Nanopore Technologies). In the sequencing process, the adaptive sampling mode was employed. In Helaly, the target sequences were set at 5,000 bp, at both ends of the HiFi contigs. In Fahl, targets were set at the first 5,000 bp, with less than 50% of repeat regions within 100,000 bp at the end of the contigs. Base calling was performed using the super-accuracy model using MinKNOW. Low-quality reads (quality value < 10) and short reads (less than 20 kb) were trimmed using Chopper software (De Coster and Rademakers, 2023). Long reads were used to scaffold the HiFi contigs using LINKS (Warren *et al*., 2015). The parameters influencing scaffolding contiguity and accuracy were tested as follows: *k*-mer length (15, 21, 31, 41, and 51-mer), a minimum number of links (5, 10, and 15), a distance between *k*-mer pairs (4,000–40,000), allowable error distance (2%, 5%, and 10%), and maximum link ratio between the two best contig pairs (0.1 and 0.3). All other parameters were set to their default values. The accuracy of the parameters was evaluated based on large discrepancies in the genome structure between cultivars using D-genies (Cabanettes and Klopp, 2018).

**Figure 1.**
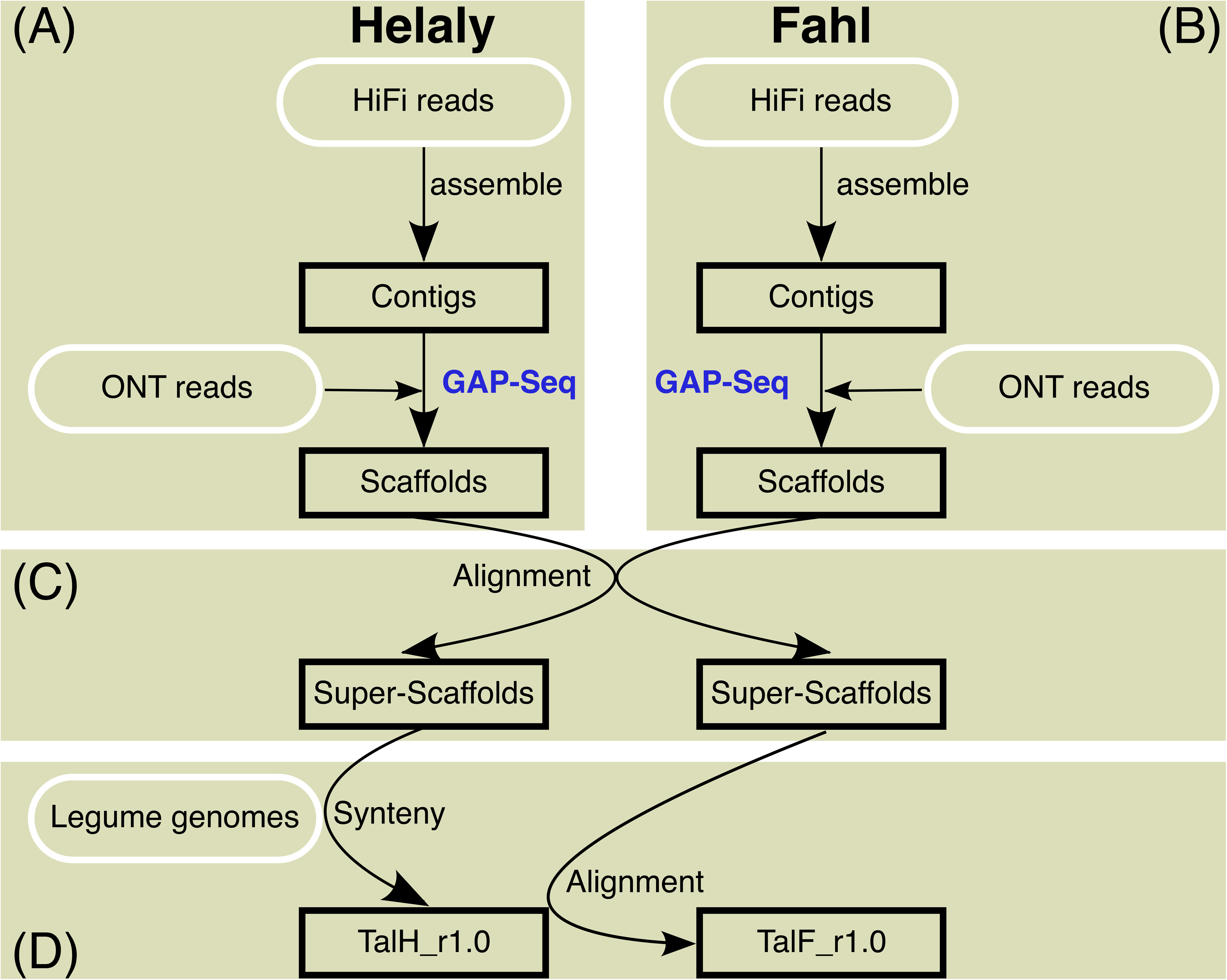
Assembly scheme of *T. alexandrinum*. The workflows and the data used for the analysis are shown in (A) for Halely and (B) for Fahl. The subsequent data analytic flows are shown in (C) and (D).

To create super-scaffolds, scaffold sequences of the two cultivars were then aligned with each other using minimap2 (v2.24) (Li, 2018) and visualized using D-genies (Cabanettes and Klopp, 2018). Based on this alignment, if one scaffold sequence from one cultivar bridged two scaffolds from the other cultivar, the latter two scaffolds were manually connected with 100 Ns to construct super-scaffolds.

Finally, based on synteny and collinearity in the legume genomes, chromosome-level sequences were generated by aligning the super-scaffolds with the chromosome sequences of the closely related species *T. repens* (Santangelo *et al*., 2023), *Medicago truncatula* (Pecrix *et al*., 2018), and *Pisum sativum* (Yang *et al*., 2022) using MCScanX (Wang *et al*., 2012). The results were visualized using SynVisio (Bandi and Gutwin, 2020). For the allotetraploid *T. repens*, because two subgenomes, probably from *T. occidentale* and *T. pallescens*, exhibit almost identical structures, only the subgenome of *T. occidentale* was used. The completeness of the genome sequences was evaluated using BUSCO v5.5.0 with embryophyte_odb10 data (Manni *et al*., 2021), LTR Assembly Index (LAI) (Ou *et al*., 2018), telomere_finder (https://github.com/MitsuhikoP/telomere_finder) (Sato *et al*., 2023), and tidk (https://github.com/tolkit/telomeric-identifier).

### Gene prediction and annotation

Potential protein-coding genes were predicted using Helixer version 0.3.1 (Stiehler *et al*., 2021), which is a gene prediction tool that combines deep learning and a hidden Markov model. To validate the results of gene prediction using Helixer, full-length cDNA sequence data were obtained using the PacBio Iso-Seq method. Total RNAs were isolated from the leaves, flowers, stems, and roots of each cultivar using the Plant Total RNA Mini Kit for Woody Plant (FAVORGEN, Taiwan) and used for library preparation according to the protocol of the ISO-Seq Express Template Preparation for Sequel and Sequel II System (PacBio). Libraries were sequenced using the Sequel II system (PacBio). Transcript isoforms for each sample were generated using the Iso-Seq analysis pipeline (PacBio) implemented in SMRT Link (PacBio). Functional annotations of predicted genes were assigned using eggNOG-Mapper 2.1.8 (Huerta-Cepas *et al*., 2017) against the eggNOG 5.0 (Huerta-Cepas *et al*., 2019) database. Gene clustering was performed using OrthoFinder (Emms and Kelly, 2019), in which the gene sequences from *Arabidopsis thaliana* (Cheng *et al*., 2017), *Glycine max* (Valliyodan *et al*., 2019), *Lotus japonicus* (Li *et al*., 2020), and *M. truncatula* (Pecrix *et al*., 2018) were involved.

Repetitive sequences were detected with RepeatMasker v4.1.6 (https://www.repeatmasker.org) using the repeat sequences obtained from the pseudomolecule sequences using RepearModeler v2.0.5 (https://www.repeatmasker.org) and from a dataset registered in Repbase (Bao *et al*., 2015).

### Population genomic analysis

Genetic variations within Helaly were investigated by double-digest restriction site-associated DNA sequencing (ddRAD-Seq) technique (Peterson *et al*., 2012). Genomic DNA was extracted from 144 Helaly seedlings using the sbeadex DNA extraction kit on the oKtopure system (LGC, UK) and used for ddRAD-Seq library construction with PstI and MspI enzymes (Shirasawa *et al*., 2016). ddRAD-Seq reads were obtained using DNBSEQ-G400 (MGI Tech) and trimmed as above. The cleaned reads were mapped onto the Helaly genome sequence as a reference using Bowtie2 (Langmead and Salzberg, 2012), and sequence variants were called using BCFtools (Danecek *et al*., 2021). High-confidence biallelic single-nucleotide polymorphisms (SNPs) were selected using VCFtools (Danecek *et al*., 2011) (parameters: minDP, 5; minQ, 200; maf, 0.05; max-maf, 0.95; and max-missing, 0.8). Linkage disequilibrium and haplotype blocks were calculated and visualized using LDBlockShow (Dong *et al*., 2020). Principal component analysis (PCA) was performed using PLINK v1.9 (Purcell *et al*., 2007). Tajima’s *D* using a sliding window approach along pseudomolecule sequences was computed using VCFTools with a non-overlapping 100-kb window size (Tajima, 1989; Danecek *et al*., 2011).

In addition to genetic variations within the cultivar, SNPs between the cultivars were detected from the short reads, which were used for genome size estimation. The cleaned reads were mapped onto the Helaly genome sequence as a reference using Bowtie2 (Langmead and Salzberg, 2012), and sequence variants were called using BCFtools (Danecek *et al*., 2021). Loci with homozygous alternative alleles in Helaly were filtered out. High-confidence biallelic SNPs were selected using VCFtools (Danecek *et al*., 2011) (parameters: minDP, 5; minQ, 200; max-missing, 1). The number of SNPs was counted using a sliding window approach of VCFTools (Danecek *et al*., 2011) with a100-kb window size.

Enrichment analyses for gene ontology (GO) were performed using topGO in the R package (Alexa and Rahnenfuhrer, 2023), and Fisher’s exact test and multiple corrections were performed with a false discovery rate.

## Results

### Genome sequence and scaffolding with GAP-Seq

On the basis of *k*-mer frequency analysis using whole genome sequences with short reads (99 Gb for Helaly and 105 Gb for Fahl), the estimated genome sizes were 486 Mb for Helaly and 474 Mb for Fahl (Figure S1), for which the peaks of *k*-mer multiplicity of 125 for Helaly and 143 for Fahl were used. The presence of two peaks in the *k*-mer distribution suggested a high level of genomic heterogeneity in both varieties.

For genome assembly of Helaly, we acquired 1.48 million HiFi reads, totaling 32.0 Gb (genome coverage of 65.8X), and assembled them into 514 contigs spanning 548 Mb with an N50 length of 14.6 Mb (Figure 1A and Table 1). To efficiently scaffold the contigs, we employed a novel strategy, i.e., gap-targeted sequencing (GAP-Seq), to target contig ends with a total length of 5.14 Mb (0.94% of the assembly size) for sequencing. Target sequencing with the Oxford nanopore technology (ONT) adaptive sampling strategy allowed sequencing of 7.69 Gb consisting of 3.35 million reads. Of these long reads, 1.28 million reads were recognized as target sequences, and 56,513 reads with sufficient accuracy (>10 QV score) and length (>20 kb) were selected as high-quality reads. More than half of the high-quality reads were omitted because of sequence stretching in the direction proximal to the target regions. Comparison of the scaffolding contiguity and accuracy among multiple parameter sets of the scaffolding software revealed that the distance between *k*-mer pairs affected the scaffolding efficiency, whereas other parameters had a small impact. To avoid mis-assembly, more conservative parameters than defaults were employed as follows: *k*-mer of 51; at least 15 links; distance between *k*-mer pairs, 20,000 bp; allowable error on this distance, 2%; and maximum link ratio between two best contig pairs, 0.1. As a result, 61 contigs were connected into 32 scaffold sequences with the GAP-Seq method. The number of resultant sequences was 482, and the contig N50 length was extended to 21.3 Mb (Figure 2A).

**Figure 2.**
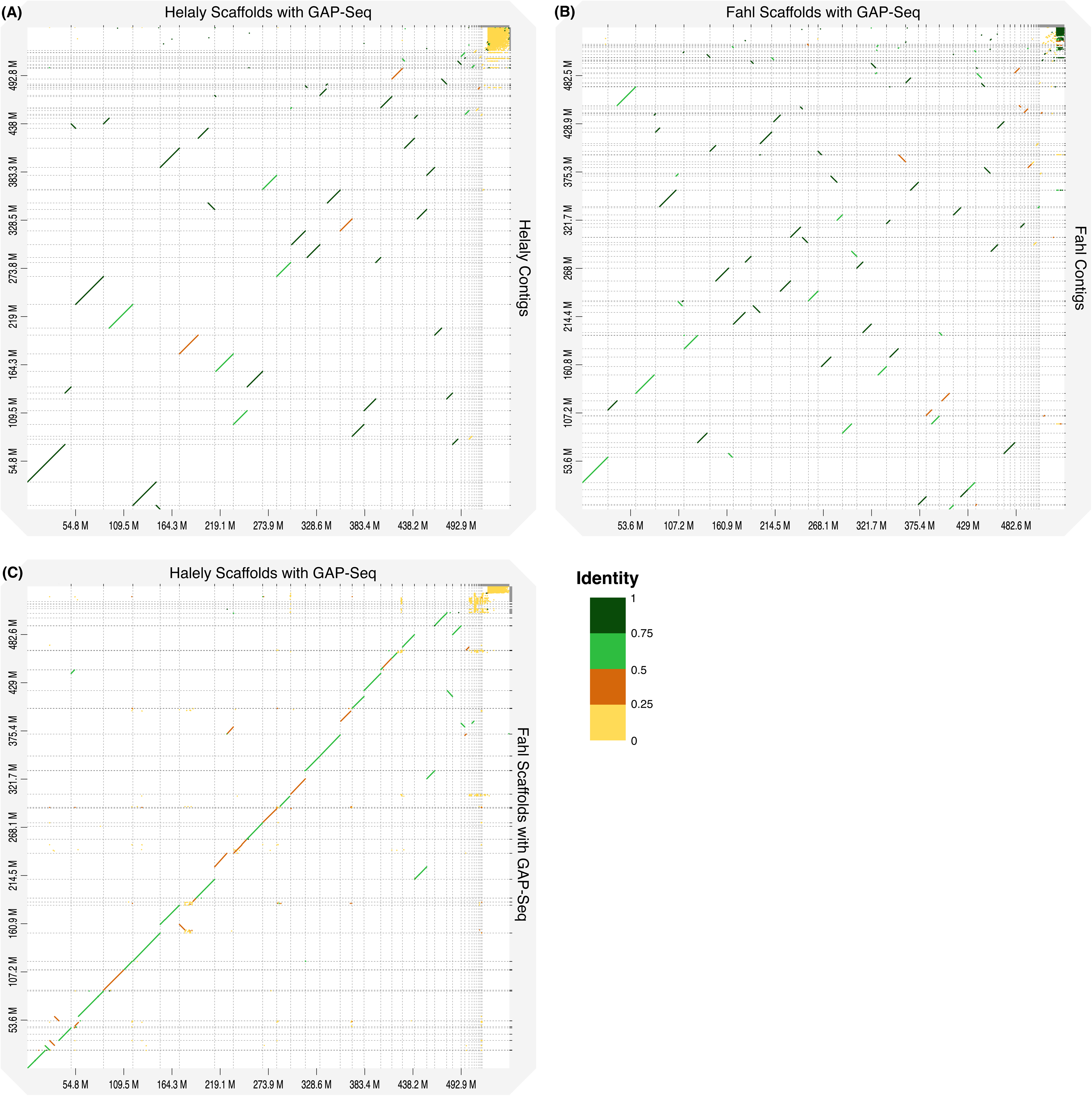
Contiguity improvement among assemblies at each step of the workflow. (A) Comparison between contigs and scaffolds using GAP-Seq in Helaly. (B) Comparison between contigs and scaffolds using GAP-Seq in Fahl. (C) Comparison of scaffolds between Helaly and Fahl. Horizontal and vertical dotted lines indicate the ranges of contigs or scaffolds. Line colors represent sequence identity.

**Table 1.** Statistics of the genome assembly and gene prediction at each step of the workflow.

|  | Contig | Scaffold | Super-scaffold | Final assembly | Pseudomolecule |
| --- | --- | --- | --- | --- | --- |
| **Helaly** |  |  |  |  |  |
| Amount of input reads (base) | 32,047,672,472 | 7,686,104,399 | - | - | - |
| Total length (bp) | 547,551,778 | 547,720,126 | 547,721,526 | 547,722,026 | 504,434,753 |
| Number of contigs | 514 | 482 | 468 | 463 | 8 |
| N50 length (bp) | 14,593,844 | 21,275,908 | 49,702,867 | 55,867,553 | 55,867,553 |
| Complete genome BUSCO | - | - | - | 99.00% | 99.00% |
| Number of genes | - | - | - | 48,862 | 39,945 |
| Complete protein BUSCO | - | - | - | 98.90% | 98.80% |
| **Fahl** |  |  |  |  |  |
| Amount of input reads (base) | 16,894,207,360 | 10,852,944,769 | - | - | - |
| Total length (bp) | 536,093,576 | 536,248,754 | 536,250,254 | 536,250,754 | 495,879,160 |
| Number of contigs | 316 | 260 | 245 | 240 | 8 |
| N50 length (bp) | 8,564,604 | 22,132,424 | 47,517,876 | 56,147,078 | 56,147,078 |
| Complete genome BUSCO | - | - | - | 99.00% | 98.80% |
| Number of genes | - | - | - | 42,966 | 39,585 |
| Complete protein BUSCO | - | - | - | 99.10% | 98.90% |

Consistent with the procedures performed for Helaly, we implemented the same strategy for Fahl. We acquired 0.81 million HiFi reads, totaling 16.9 Gb (genome coverage of 35.6X), and assembled them into 316 contigs spanning 536 Mb with an N50 length of 8.56 Mb (Figure 1B, Table 1). Using the GAP-Seq strategy for 915-kb (0.17%) target regions, 10.9 Gb consisting of 8.28 million ONT long reads were obtained. Of these ONT reads, 829 thousand reads were recognized as target sequences and 72,353 reads had sufficient accuracy (>10 QV score) and length (>20 kb). With the high-quality ONT reads, 79 contigs were connected and generated 260 scaffolds. The N50 length was extended to 22.1 Mb (Figure 2B).

### Constructing near-telomere-to-telomere genome assemblies

The scaffolds were manually integrated into super-scaffolds based on the alignments between the Helaly and Fahl cultivars (Figure 1C, 2C). In total, 482 Helaly scaffolds and 260 Fahl scaffolds were complementary aligned to obtain 13 super-scaffolds each for Helaly and Fahl, all of which were longer than 10 Mb. Other short scaffolds that were not aligned were retained as unplaced scaffolds of the *T. alexandrinum* genome sequences. The N50 lengths improved to 49.7 Mb and 47.5 Mb for Halely and Fahl, respectively.

To construct chromosome-level assemblies, the 13 super-scaffolds of Helaly were aligned with a subgenome of *T. occidentale* haplotype sequence of *T. repens*, based on synteny and collinearity (Figure 1D, Figure 3). Five super-scaffolds were consistent with the five chromosomes of the *T. occidentale* haplotype sequence. The other eight super-scaffolds were aligned to the remaining three *T. occidentale* chromosomes to generate three pseudomolecule sequences. These processes yielded eight pseudomolecule sequences corresponding to the basic number of *T. alexandrinum* chromosomes. Subsequently, the 13 super-scaffolds of Fahl were aligned with Helaly, in which five super-scaffolds corresponded to five chromosomes and the remaining eight super-scaffolds were connected to three pseudomolecule sequences. In both varieties, the resultant assemblies, which included both the pseudomolecule sequences and the retained short scaffolds, were named TalH_r1.0 for Helaly (547.7 Mb in total length) and TalF_r1.0 for Fahl (536.3 Mb). The nomenclature and direction of the pseudomolecule sequences were based on the eight chromosomes of the *T. occidentale* haplotype sequence of the *T. repens* genome.

**Figure 3.**
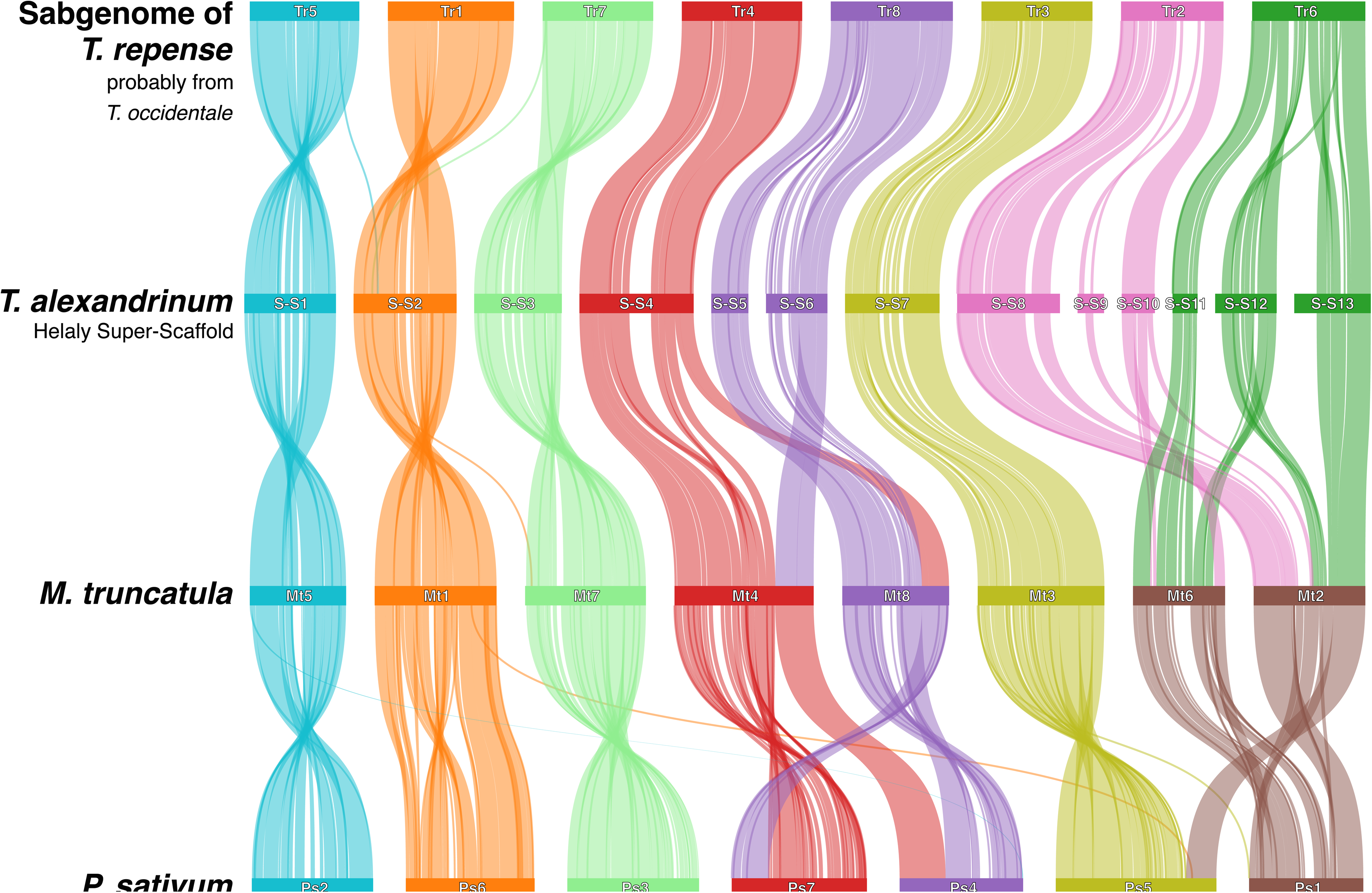
Synteny and collinearity of genes in *T. repens*, *T. alexandrinum* (Halely, super-scaffold), *M. truncatula*, and *P. sativum*. A subgenome, probable *T. occidentale* was used to represent *T. repens*.

The quality of the pseudomolecule sequences was assessed using three methods. In both TalH_r1.0 and TalF_r1.0, telomere repeats were identified at all ends of the eight pseudomolecule sequences (Supplementally Figure S2). The complete BUSCO score of TalH_r1.0 was 99.0%, and the single-copy and duplicated BUSCO scores were 94.5% and 4.5%, respectively. The complete BUSCO score of TalF_r1.0 also was 99.0%, and the single-copy and duplicated BUSCO scores were 94.1% and 4.9%, respectively. LAI scores were 21.0 and 20.7 for TalH_r1.0 and TalF_r1.0, respectively.

### Gene prediction and repeat sequence analysis

A total of 48,862 and 42,966 potential protein-coding sequences were identified based on *ab initio* prediction using Helixer in TalH_r1.0 and TalF_r1.0, respectively. Of these potential protein-coding sequences, 39,945 and 39,585 genes were located on the eight pseudomolecule sequences (Table 1). The complete BUSCO score of the predicted genes in TalH_r1.0 was 98.9%, and the single-copy and duplicate BUSCO scores were 93.4% and 5.5%, respectively. The BUSCO score of the predicted genes in TalF_r1.0 was 99.1%, and the single-copy and duplicate BUSCO scores were 93.2% and 5.9%, respectively.

The accuracy of the predicted genes was assessed using Iso-Seq data from four tissues: leaves, flowers, stems, and roots. A total of 1.52 million reads were clustered into 71 thousands (k) (leaf), 104 k (flower), 100 k (stem), and 9 k (root) full-length transcripts, and the transcripts were further clustered into 166 k unique transcripts. Similarly, Fahl exhibited full-length transcript counts of 77 k (leaf), 79 k (flower), 91 k (stem), and 5 k (root), totaling 150 k unique transcripts. The complete BUSCO score of the unique transcripts was 94.6% for Helaly and 92.6% for Fahl. The predicted gene models with Helixer matched 145,410 (97.2%) and 135,440 (97.6%) of the unique transcripts for Helaly and Fahl, respectively, indicating that the accuracy of the predicted genes was well supported by the transcriptome data.

Repeat sequences occupied 354.2 Mb (64.67%) and 342.9 Mb (63.93%) in TalH_r1.0 and TalF_r1.0, respectively (Table S1). The dominant repetitive sequences were long terminal repeat (LTR) elements, constituting 27.79% of the sequences in Helaly and 29.55% in Fahl, followed by unclassified repeats, which constituted 14.64% of the sequences in Halely and 15.94% in Fahl. Retroelements and unclassified repeats were fewer in Helaly than in Fahl, whereas DNA transposons were more numerous in Helay than in Fahl (10.89% and 6.28%, respectively). The differences in repeat sequences between Helaly and Fahl included *hobo-Activator* elements, which were 15 Mb larger in Helaly (three times more), primarily on chromosomes 2 (20 times more) and 4 (39 times more). MULE-MuDR elements were 5.5 Mb larger in Helaly (1.6 times more) across all chromosomes. In contrast, LTR elements were 6.3 Mb larger in Fahl (1.04 times more) across all chromosomes except chromosome 6.

### Ortholog analysis of the *T. alexandrinum* genes

For further evaluation of gene prediction accuracy, we conducted an ortholog analysis focusing on orthogroup sharing and phylogenetic relationships, confirming evolutionary conservation, and identifying lineage-specific gene acquisition. The genes predicted in TalH_r1.0, which were used as representatives of *T. alexandrinum*, were clustered with those of three legume species, *G. max*, *L. japonicus*, and *M. truncatula*, as well as *A. thaliana* (Figure 4). In total, 25,008 clusters were identified across the five species. Of these, 12,406 (49.6%) were shared among the five species, and 2,975 (11.9%) were shared among the four legume species. Species-specific clusters were identified, with 1,114 (4.45%) clusters found in *A. thaliana*, 1,363 (5.45%) in *G. max*, 391 (1.56%) in *L. japonicus*, 1,138 (4.55%) in *M. truncatula*, and 1,421 (5.68%) in *T. alexandrinum*. The number of clusters shared exclusively with *T. alexandrinum* was 16 (0.06%) for *A. thaliana*, 162 (0.65%) for *G. max*, 64 (0.26%) for *L. japonicus*, and 1,278 (5.11%) for *M. truncatula*.

**Figure 4.**
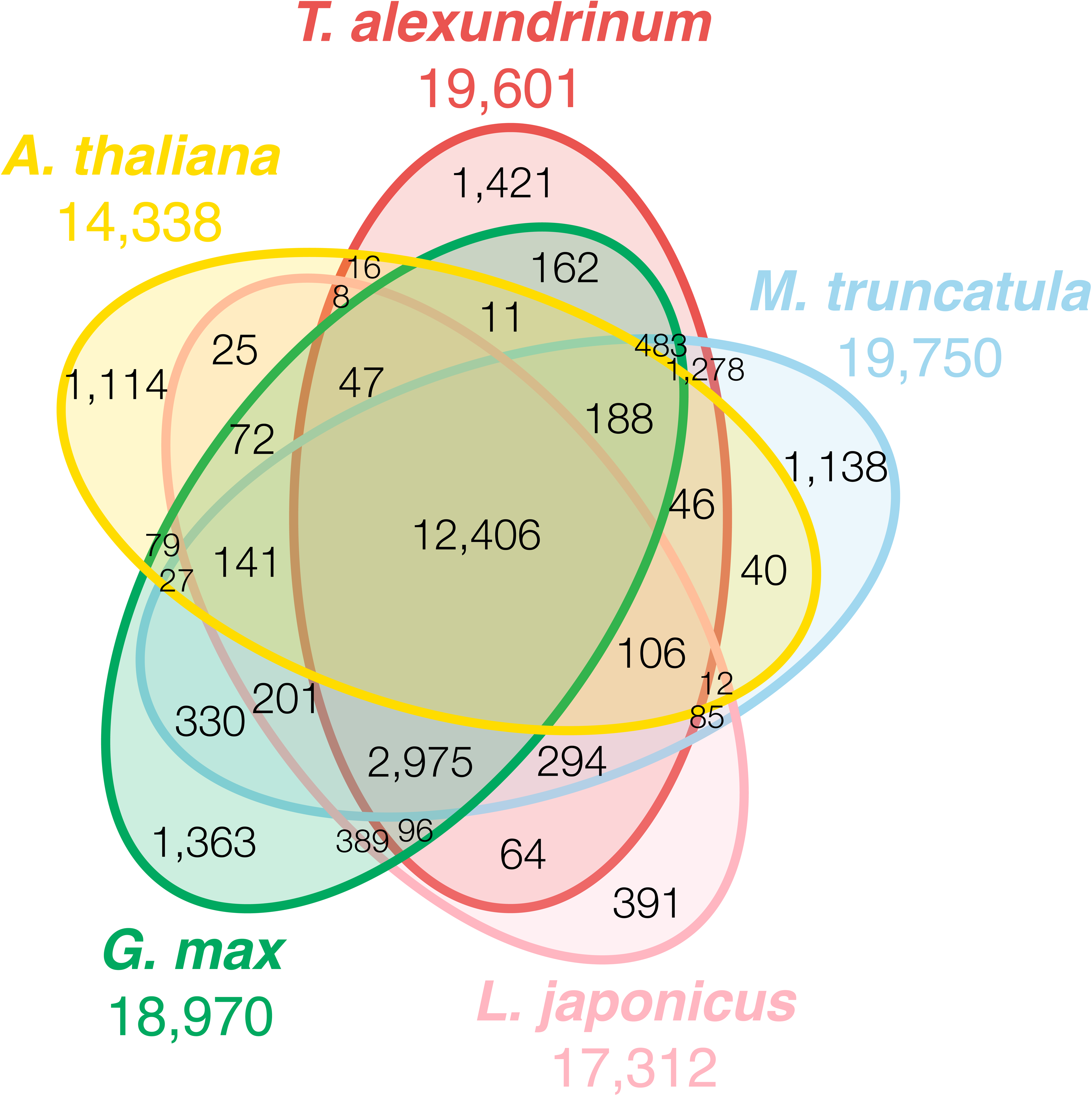
Venn diagram of the numbers of gene clusters in the four legume genomes and *A. thaliana*.

### Population genomic analysis and GO enrichment analysis

To evaluate the genetic diversity within and between Egyptian clover cultivars, a genome-wide SNP analysis was performed. First, to evaluate genetic diversity level within a cultivar, we obtained 958 million ddRAD-Seq reads from 144 Helaly individuals. The reads were mapped on the TalH_r1.0 genome sequence with a mapping rate of 75.6% (61.1%- 86.9%) to detect 17,031 SNPs. Then, Tajima’s *D* was calculated for the whole genome using 100-kb sliding windows. The average Tajima’s *D* was 1.82 (Figure 5), which value was evenly distributed over the genome.

**Figure 5.**
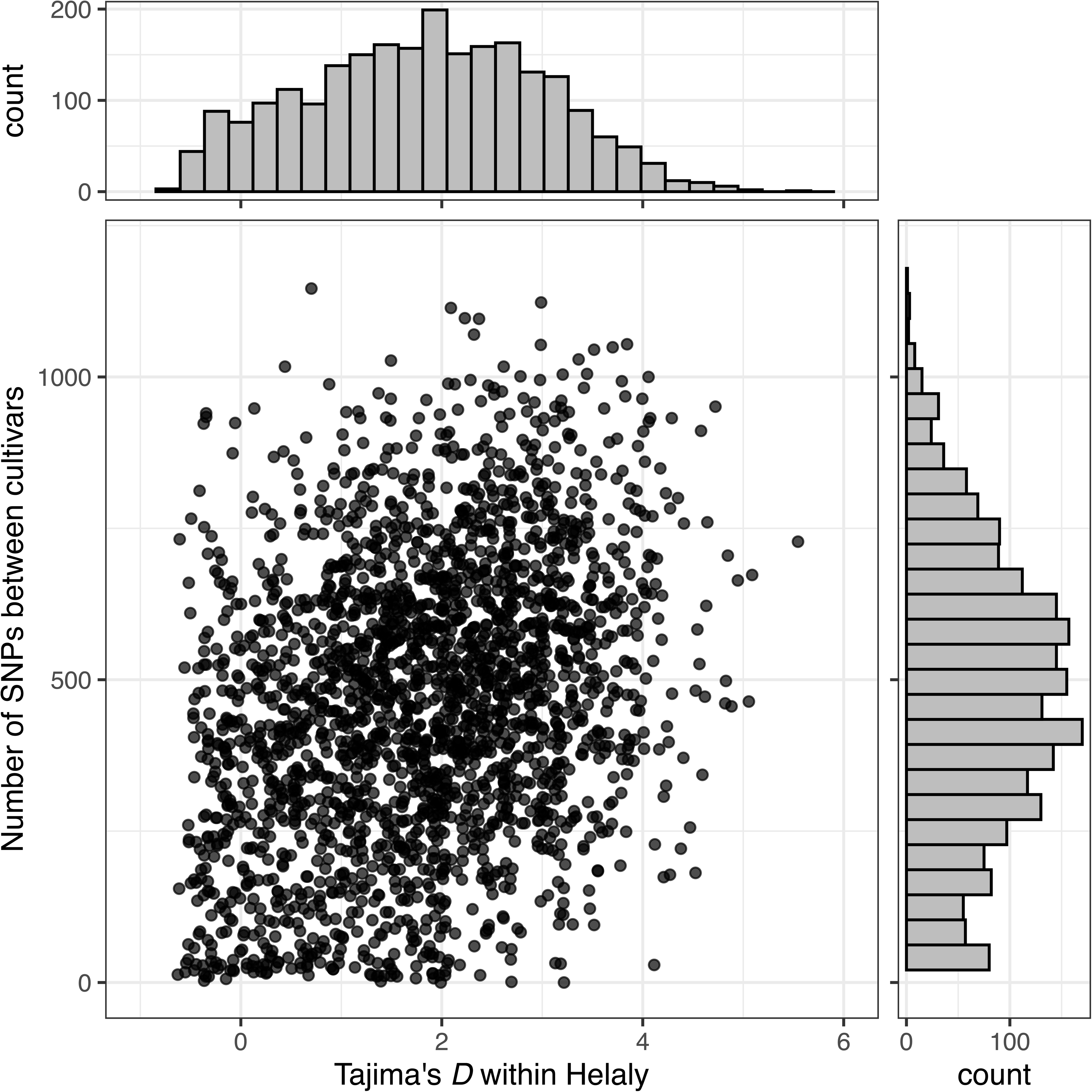
Relationships between Tajima’s *D* within Helaly and the number of SNPs between cultivars for each 100 kb sliding window of genomic regions. The bar plots at the top and right display the histograms of Tajima’s *D* and the number of SNPs, respectively.

Second, to investigate genetic diversity level between cultivars, the short reads for Helaly and Fahl were mapped onto the Helaly genome sequence as a reference to detect 1,677,056 SNPs between the two varieties. We focused on 38 SNP-rich regions with low Tajima’s *D* values because these regions exhibited high genetic diversity levels between the two cultivars but low levels in Helaly (Figure 6). Indeed, a slight positive correlation was observed between the number of SNPs between the cultivars and Tajima’s *D* within Helaly (correlation coefficient = 0.3, p-value = 6.95 x 10-49).

**Figure 6.**
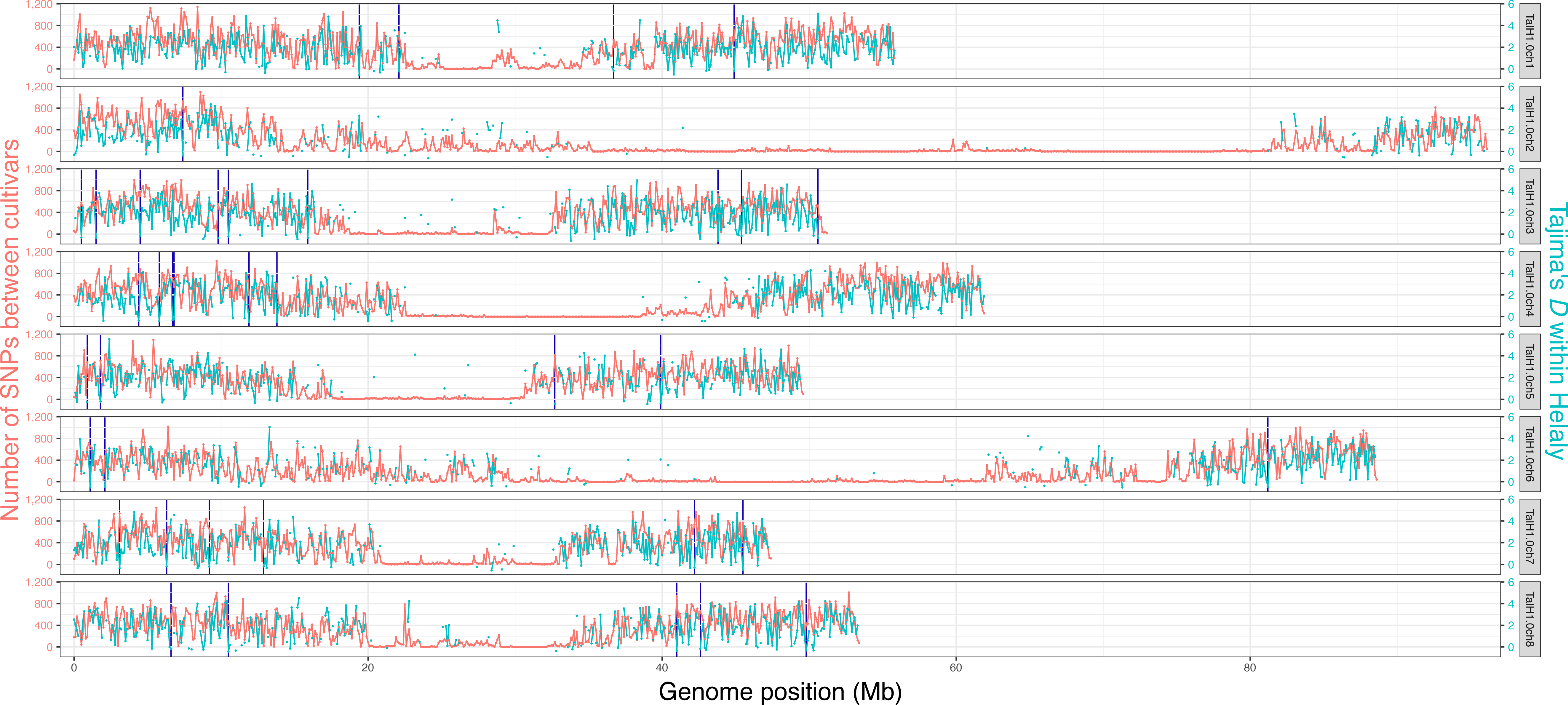
Tajima’s *D* within Helaly and the number of SNPs between cultivars along each chromosome with 100 kb sliding windows. Blue lines indicated the 38 genomic regions that might determine the unique characteristics of Helaly and identified in this study.

We performed GO enrichment analysis of 453 genes in the 38 genomic regions against all genes of Helaly. Forty GO terms were significantly enriched (p < 0.01), and 11 terms were enriched with multiple corrections (q-value < 0.01; Table S3). The enriched GO terms included cell wall and structural processes, metabolic and biosynthetic processes, stress response and defense, development, chromatin and transcription processes, apoptosis, and aging.

## Discussion

We constructed a near-T2T genome assembly consisting of eight pseudomolecule sequences corresponding to the basic number of chromosomes in two Egyptian clover cultivars. The total assembly sizes of Helaly and Fahl were 547.7 Mb and 536.3 Mb, respectively. These sizes were larger than the estimated sizes for Helaly (486 Mb) and Fahl (474 Mb). High heterozygosity and repetitive sequences might have been responsible for the inconsistency between an assembly size and an estimated size based on *k*-mer frequency analysis (Liu *et al*., 2013). The underestimation observed in the present study may be due to the high heterozygosity and the abundance of repetitive sequences in the *T. alexandrinum* genome.

The assembly size for Helaly (547.7 Mb) was 11.4 Mb larger than that for Fahl (536.3 Mb), reflecting a size difference of 11.3 Mb in repeat sequences between the two cultivars: 354.2 Mb in Helaly and 342.9 Mb in Fahl. This difference in the size of the repeat sequences can be primarily attributed to the expansion of certain DNA transposons, *hobo*-*Activator* and MULE-MuDR, in the Helaly genome. Both transposable elements are widespread among angiosperms and induce mutations that enhance genomic studies on maize (Ros and Kunze, 2001; Lisch, 2015). Furthermore, the transposable elements contribute to breeding by creating genetic diversity and enabling the selection of beneficial mutations (Lisch, 2013).

Filling sequences in gap regions consisting undetermined nucleotide sequences and orientation of small contigs in correct direction are essential to produce reliable genome assemblies. To achieve the near-T2T genome assemblies, we developed a novel scaffolding strategy, GAP-Seq, which enables to determine nucleotide sequences in gap regions by a target sequencing technology based on the adaptive sampling method (ONT). Using the GAP-Seq approach, 61 contigs were connected in Helaly, improving the N50 length from 14.6 Mb to 21.3 Mb. In Fahl, 79 contigs were connected, with the N50 length improving from 8.56 Mb to 22.1 Mb. This method has an advantage over the practical target enrichment technologies requiring molecular experiments using PCR or probes and labor costs, because the adaptive sampling method can computationally enrich the targets on the flow cell of the ONT sequencer without the costs. PacBio HiFi together with GAP-Seq based on ONT adaptive sampling can be applicable for T2T genome assembly in other plant species.

Egyptian clover cultivars are not pure lines because of their allogamy, to avoid inbreeding depression. As reported in many previous studies (Moeller *et al*., 2007; Mulugeta *et al*., 2023), cultivar-specific characteristics can be identified using the population genomic analysis to examine genetic variations within and between cultivars. In this study, we evaluated the genetic diversity level within Helaly. Since the positive Tajima’s *D* value was evenly distributed over the genome, a particular population structure or dynamics was supposed in this cultivar. Therefore, the Helaly is suggested to be a composite population of subgroups. This result is consistent with the fact that Helaly was bred using a mass selection method with pollinators, a common practice in forage breeding programs. Furthermore, we identified 38 genomic regions under relatively strong purifying selection that differed between the cultivars. Indeed, Helaly-specific characteristics, such as high productivity and stress resistance, were supported by GO enrichment analysis, in which two categories were enriched: growth and response to stress (Table S3). These results indicate that the characteristics of Helaly might be derived from at least 38 genomic regions and that the difference between the two cultivars might be explained by the GO terms enriched in the regions. Cultivar-specific loci could confer desirable traits during breeding. The identification of genetic loci for agriculturally important traits can facilitate Egyptian clover breeding programs with marker-assisted selection through genome-wide association studies and quantitative trait loci mapping.

In this study, we establisehd the genome assemblies of two Egyptian clover cultivars to reveal that the genome size and structure differed between the two cultivars. These data suggest that specific genomic sequences are present in cultivars and individuals. Therefore, high-quality genome sequences across cultivars and individuals, known as pan-genomics, are required to understand the genetic variations in the cultivars and individuals of a species.

## Acknowledgments

We thank K. Ozawa, C. Minami, H. Tsuruoka, Y. Kishida, and A. Watanabe (Kazusa DNA Research Institute) for their technical assistance.

## Data availability

Raw sequencing reads and assemblies were deposited in the DNA Data Bank of Japan (DDBJ) under the accession number PRJDB818306. Genomic information is available from Plant GARDEN (https://plantgarden.jp/).

## Supporting Data

Supporting Figure S1. Estimation of the genome size of *T. alexandrinum*, based on *k*-mer analysis (*k* = 21) with given multiplicity values. Triangles indicate the peak used by estimation.

Supporting Figure S2. The distribution of telomere repeats along eight chromosomes for Helaly and Fahl. Colors indicate each chromosome.

Supporting Table S1. Repetitive sequences in TalH_r1.0 and TalF_r1.0.

Supporting Table S2. Number of reads and aaRAD-Seq mapping rates.

Supporting Table S3. Enriched gene ontology list of Helaly-specific genomic regions.

