## Supplementary figures and images for "Near-complete telomere-to-telomere *de novo* genome assemblies of Egyptian clover (*Trifolium alexandrinum*)"

### Supplementary Figure S1

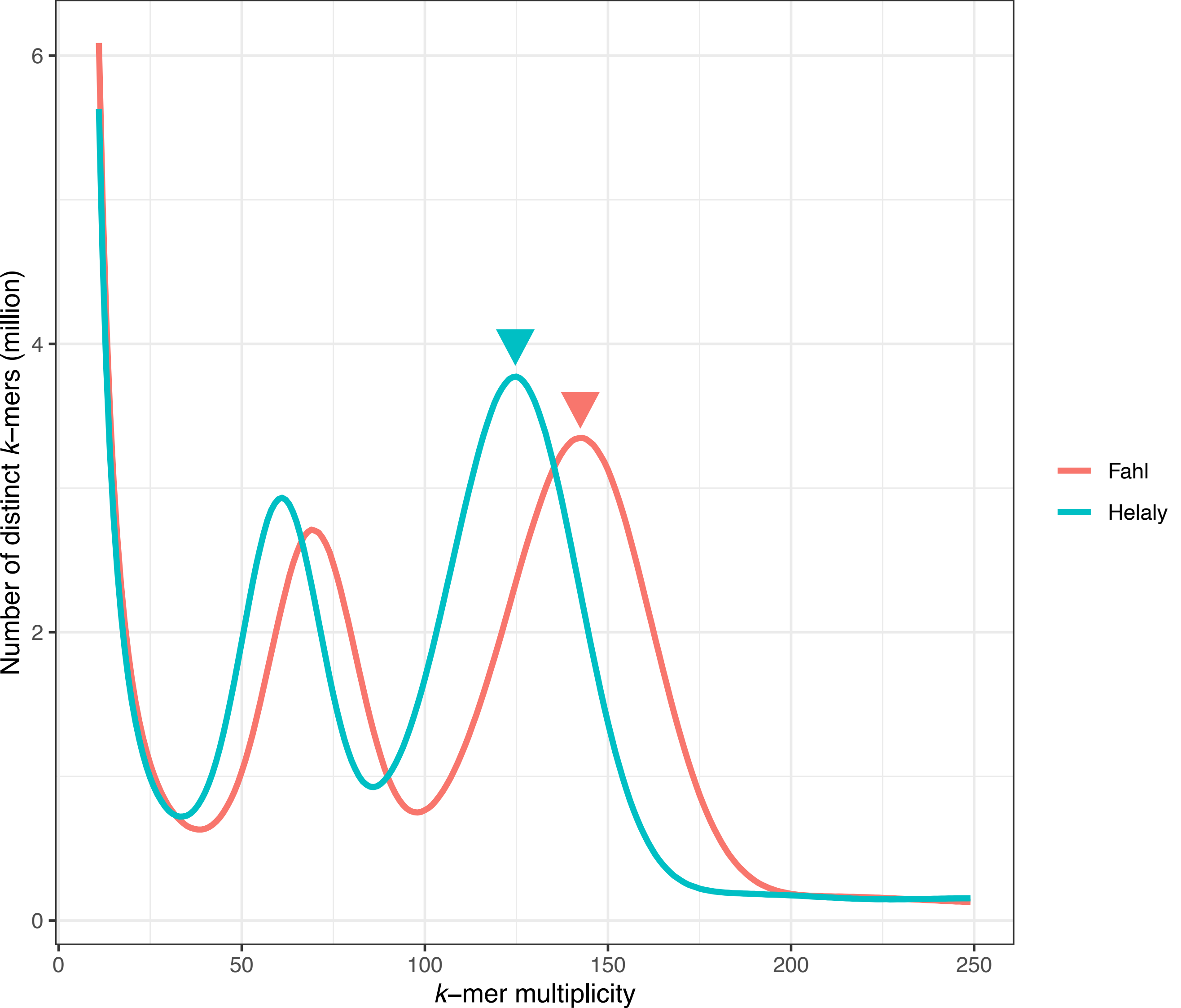

### Supplementary Figure S2

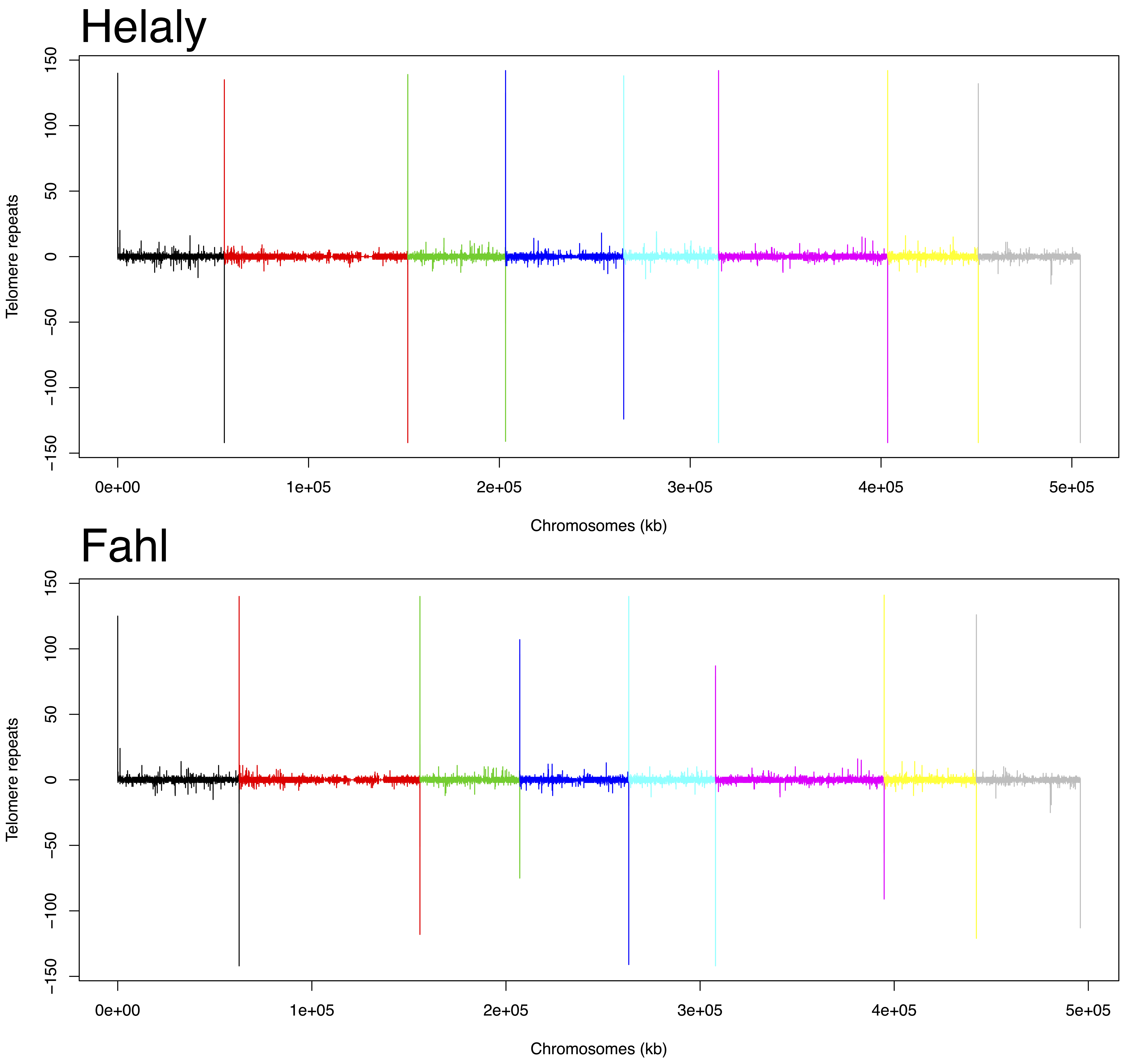
